# High spatial resolution analysis using indentation mapping differentiates biomechanical properties of normal vs. degenerated mouse articular cartilage

**DOI:** 10.1101/2021.10.26.465857

**Authors:** Anand O. Masson, Bryce A. Besler, W. Brent Edwards, Roman J Krawetz

## Abstract

Characterizing the biomechanical properties of articular cartilage is crucial to understanding processes of tissue homeostasis vs. degeneration. In mouse models, however, limitations are imposed by their small joint size and thin cartilage surfaces. Here we present a 3D automated surface mapping system and methodology that allows for mechanical characterization of mouse cartilage with high spatial resolution. We performed repeated indentation mappings, followed by cartilage thickness measurement via needle probing, at 31 predefined positions distributed over the medial and lateral femoral condyles of healthy mice. High-resolution 3D x-ray microscopy (XRM) imaging was used to validate tissue thickness measurements. The automated indentation mapping was reproducible, and needle probing yielded cartilage thicknesses comparable to XRM imaging. When comparing healthy vs. degenerated cartilage, topographical variations in biomechanics were identified, with altered thickness and stiffness (instantaneous modulus) across condyles and within anteroposterior sub-regions. This quantitative technique comprehensively characterized cartilage function in mice femoral condyle cartilage. Hence, it has the potential to improve our understanding of tissue structure-function interplay in mouse models of repair and disease.

## Introduction

The articular cartilage in synovial joints has a specialized three-dimensional structure and biochemical composition that provides low-friction, wear-resistance, and load-bearing properties to the tissue. Mechanical loading is essential to tissue homeostasis and influences gene expression, chondrocyte metabolism, extracellular matrix (ECM) maintenance, and associated interstitial fluid permeability ^1,2^. Therefore, the biomechanical properties of cartilage are unique indicators of tissue homeostasis versus degeneration. Indeed, research into cartilage-related changes during the development of degenerative diseases such as osteoarthritis (OA) has demonstrated that early tissue dysfunction involves loss of proteoglycans and increased water content, resulting in reduced compressive strength and higher tissue permeability ^3,4^. Disease progression leads to further structural damage and altered mechanics ^5^, ultimately resulting in tissue failure.

Mouse models are commonly used in pre-clinical studies focused on cartilage repair and pathophysiology of OA ^6^. Mice are advantageous because of the well-established development of genetically modified strains ^7^ and the availability of diverse model systems mimicking mechanisms of spontaneous and induced tissue degeneration and regeneration ^8–11^. In this regard, mouse models serve as powerful tools for the targeted assessment of cellular and molecular processes and the discovery of novel therapeutics related to cartilage regeneration and OA. Yet, while the structural integrity and biochemical composition of murine cartilage are routinely assessed through histological and molecular approaches, the evaluation of how these features translate into mechanical function is limited. The main challenge in mechanical function assessment stems from their small joint size and thin cartilage found in mice relative to other species ^12^. Prior efforts to overcome these challenges include finite element modelling and optimization of small-scale indentation techniques ^3,13–15^.

Mechanical indentation (including AFM techniques), performed using either creep or stress-relaxation protocols, is widely employed for assessing the biomechanical behavior of cartilage and OA-related changes in many species ^16–18^, and considered the gold standard for small animal joints ^19^. Unlike confined and unconfined compression tests, no sectioning or subsampling of tissue (i.e., cylindrical explants) is required ^20^. Instead, the cartilage tissue and its subchondral bone interface is kept intact, providing more physiologically relevant data. Indentation is also advantageous in that it is non-destructive and allows for repeated measures *in situ* ^21^. However, the natural curvature of joint articular surfaces poses a challenge to testing, as indentation must be conducted perpendicularly to the surface ^22^.

Recently, a novel automated indentation technique has been developed for the mechanical assessment of cartilage ^23^. This commercially available multi-axial apparatus (Mach-1™, *Biomomentum Inc*, Laval, QC) is capable of detecting specimen surface orientation at each position of measurement and subsequently indent normal to the surface ^23,24^. As such, it can map entire cartilage surfaces using a single setup with high spatial density, and has been previously shown to discriminate between healthy and diseased human cartilage samples ^23,25^. This apparatus has been recently employed by Woods and colleagues ^26^ to evaluate altered biomechanics in a mouse model of cartilage degeneration. However, the study was conducted in the non-load-bearing region of the mouse knee joint. Considering that the knee range of motion in mice is between 40° to 145° (i.e., unable to fully extend) ^27^, with normal gait range between 90.5° and 120° (extension-flexion) ^28^, the contact regions in the distal femoral condyles are located further posteriorly compared to humans. Careful consideration of species-specific differences in knee joint anatomy and kinematics is imperative for proper translation of pre-clinical models ^29^.

Woods and colleagues ^26^ were also unable to account for site-specific cartilage thickness variations in their measurements, instead using the mean cartilage thickness obtained via histological analysis in their assessment ^26^. Other studies have employed mean cartilage thickness values retrieved through histology or imaging techniques to characterize and model tissue mechanical parameters in mice ^13,15^. Implicit in this approach is the underlying assumption that cartilage thickness is relatively uniform among medial and lateral compartments (e.g., femoral condyles or tibial plateaus) and along their anteroposterior or mediolateral axis. Yet, regional variations in thickness are recognized within these cartilage surfaces ^12,15,30^, and can impact indentation measurements ^22,31^.

Hence, the purpose of this study was to investigate the reliability of automated indentation mapping in the assessment of healthy femoral articular cartilage in mice and characterize site-specific variations in cartilage thickness. We also employed concurrent contrast-enhanced 3D x-ray microscopy (XRM) imaging to validate the cartilage thickness measurements from our optimized needle probing protocol^32^. Finally, this approach was used to investigate biomechanical differences in a clinically relevant mouse model of cartilage degeneration ^33^. Together, we show that automated indentation is reliable and able to characterize topographic and mechanical variations across condyle cartilage locations in intact cartilage. Moreover, this technique was able to identify regional changes in cartilage thickness and stiffness in degenerated cartilage. A comprehensive and standardized biomechanical evaluation of cartilage in repair and disease can greatly contribute to our understanding of tissue structure-function interplay, thereby enhancing the clinical relevance of mouse models.

## Results

### Automated indentation mapping reliability

While previous studies have employed indentation mapping on mouse cartilage, to our knowledge none have measured its accuracy and precision. Therefore, we performed three repeated mappings per position for all specimens to undertake a reliability analysis of the automated surface indentation technique. Each of the 31 predefined positions distributed over the femoral condyles of ten C57Bl6 mice was included in this analysis (Figure 1A). The setup was developed and optimized (Figure S1) to assess the load-bearing regions of the femoral condyles and achieve non-destructive retrieval of specimen post-testing for subsequent 3D XRM imaging analysis. The imposed step deformation (i.e., indentation depth) on the femoral cartilage yielded typical stress-relaxation behavior, characterized by a sharp increase in force followed by gradual relaxation over time until equilibrium (Figure 1B). Assessment of stress-relaxation and corresponding force-displacement curves (Figure 1B) demonstrated consistency among repeated measurements for single positions and visible differences in peak reaction forces between condyles. These observations were further evidenced by the spatial distribution of peak force values across condylar testing sites (Figure 1C, Table S1). A total of 930 indentation measurements were retrieved, out of which only 21 produced invalid curves (seven testing sites at specimen’s periphery with higher angles yielded noisy signals, Figure 1C), representing a 2.26 % error rate during data acquisition. High reliability and absolute agreement between repeated measures for individual testing sites were observed, with 4.7% intra-assay average coefficient of variation (CV) (Table S1) and intraclass correlation coefficients - ICC (lower 95%, upper 95%) - ranging from 0.974 (0.966, 0.981) for the lateral condyle to 0.971 (0.963, 0.978) for the medial condyle. Mean peak force values illustrate site-specific variations within and between condyles (Figure 1D). The lateral condyle values varied significantly per position (*p <* 0.0001), ranging from 0.07 to 0.15 N and showed a trend for higher values at outermost positions, with a slight decrease in force posteriorly. The latter was also seen for the medial condyle, wherein heterogeneities in peak force were also apparent (*p <* 0.0001) and had a wider range – from 0.15 to 0.294 N. Since the analysis per testing site also reflects inherent deviations due to anatomical positioning across specimens, data was pooled for regional (between condyles) and sub-regional (between and within anteroposterior locations) comparisons.

**Figure 1.**
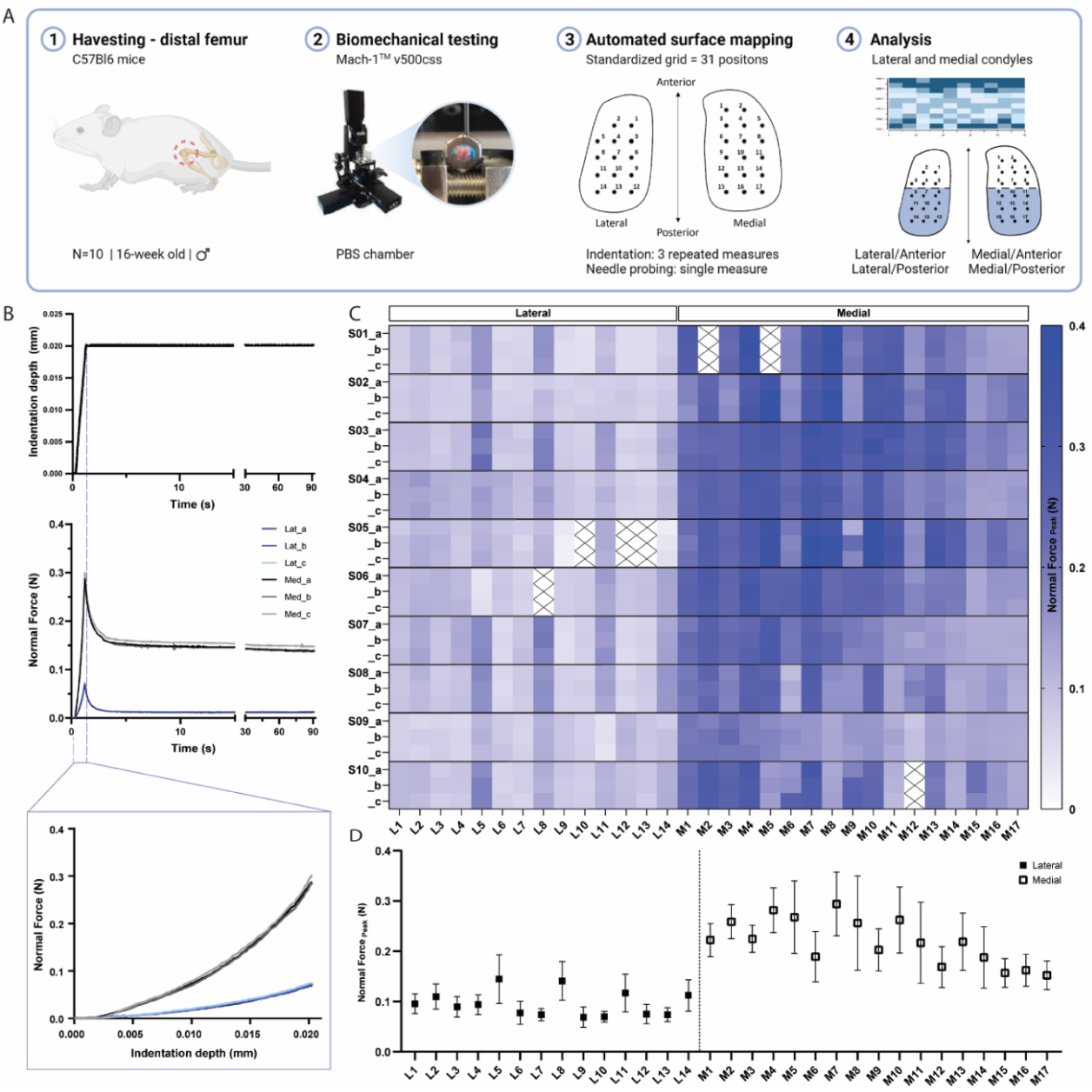
(A) Schematic overview of experimental design employed for biomechanical testing of murine articular cartilage using Mach-1™ v500css mechanical tester (B) Step displacement used for cartilage indentation (top), with typical force-relaxation response curves obtained for three repeated measures on representative lateral and medial condyle positions (middle) and corresponding force increase with indentation depth (bottom) (C) Normal peak force recorded for all three repeated measures (a-c) considering each of the 31 testing sites, L1-L14 at lateral condyle and M1-M17 at medial condyle, for each specimen (*n =* 10, S01-S10), demonstrates general agreement for intra-specimen measurements on both condyles (D) Mean peak force values varied within and between condyle locations and higher within medial condyle testing sites. Data is presented as mean ± SD.

As Table 1 shows, the average peak force was significantly higher on the medial condyle and on both its anterior and posterior sub-regions when compared to lateral counterparts (Lat/Ant vs. Med/Ant and Lat/Post vs. Med/Post). Interestingly, no significant differences were observed between sub-regions of the lateral condyle (Lat/Ant vs. Lat/Post). In contrast, the mean peak force yielded at the Med/Post sub-region was 20% lower than on the Med/Ant (*p <* 0.01). As cartilage thickness variations between and within condyle locations could affect peak forces measured at same indentation depth ^34^, with thinner cartilage yielding higher force values, we sought to determine the cartilage thickness distribution within the same surfaces and validate this approach using XRM imaging.

**Table 1.**
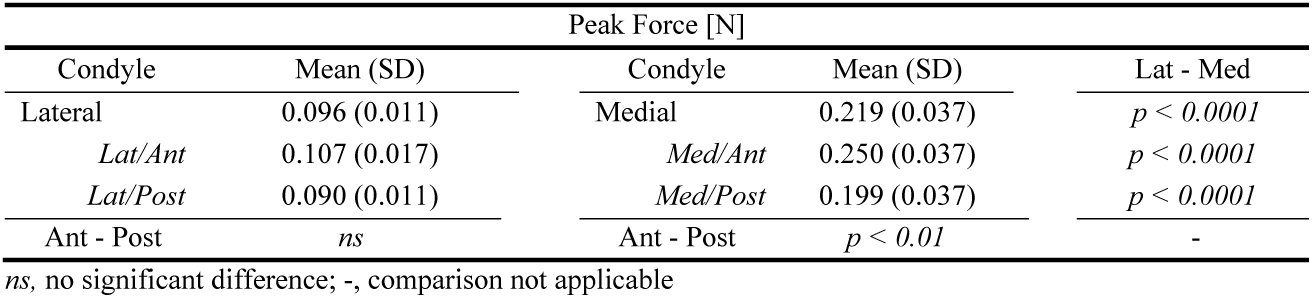
Mean and standard deviation (SD) values for peak force (N) as determined by automated indentation test performed for *n* = 10 distal femur samples of murine AC. Mean values compared between condyles (Lateral/Medial; unpaired, Student’s *t* test, *α* = 0.05) and within sub-regions of condyles (Lat/Ant, Lat/Post, Med/Ant, Med/Post; *one-way* ANOVA, *p* < 0.05).

### Cartilage thickness characterization: comparison between needle probing and XRM imaging

Needle probing (NP) thickness mapping was performed on all ten femoral condyles, which were subsequently scanned using contrast enhanced XRM imaging. As shown in Figure 2A, reconstructed 3D datasets of murine distal femurs allowed us to validate the spatial distribution of NP testing sites, thereby the corresponding ROI coordinates for imaging processing could be determined. Additionally, the 2D slices confirmed that the needle probe pierced the full length of the cartilage, reaching the subchondral bone (Figure 2A). The cartilage surface and cartilage-subchondral bone interface positions were identified using the load-displacement curves from NP (Figure 2B) and the cartilage thickness for each position was then calculated considering the surface angle (*Methods*). There was a 2.58% rate of needle probe failure (8 out of 310 measurements, testing sites near the specimen’s edge) during data acquisition. NP and contrast enhanced XRM imaging yielded similar cartilage thickness distributions on both condyles (Figure 2C), demonstrating a highly significant correlation between paired values (R = 0.842, *n* = 302, *p <* 0.0001) (Figure 2D). Further method-comparison using a Bland-Altman plot (Figure 2E) illustrated XRM measurements were approximately 6.8 µm thicker on average than needle probing, representing only a 1.55 voxels difference (4.39 µm resolution - XRM). Moreover, no significant differences were found between pairwise mean thickness values (NP vs. XRM) for individual positions within the lateral condyle (Figure 2F). Similar results were shown for the medial condyle, wherein higher mean thickness values from XRM were only seen on M06 (Δ = 8.68 µm, *p =* 0.028), M11 (Δ = 9.89 µm, *p* < 0.01), and M13 (Δ = 9.34 µm, *p =* 0.012) testing sites. No obvious reason for these localized differences were found, and dullness of needle was ruled out since subsequent testing sites (M14 to M17) yielded comparable cartilage thickness values between techniques.

**Figure 2.**
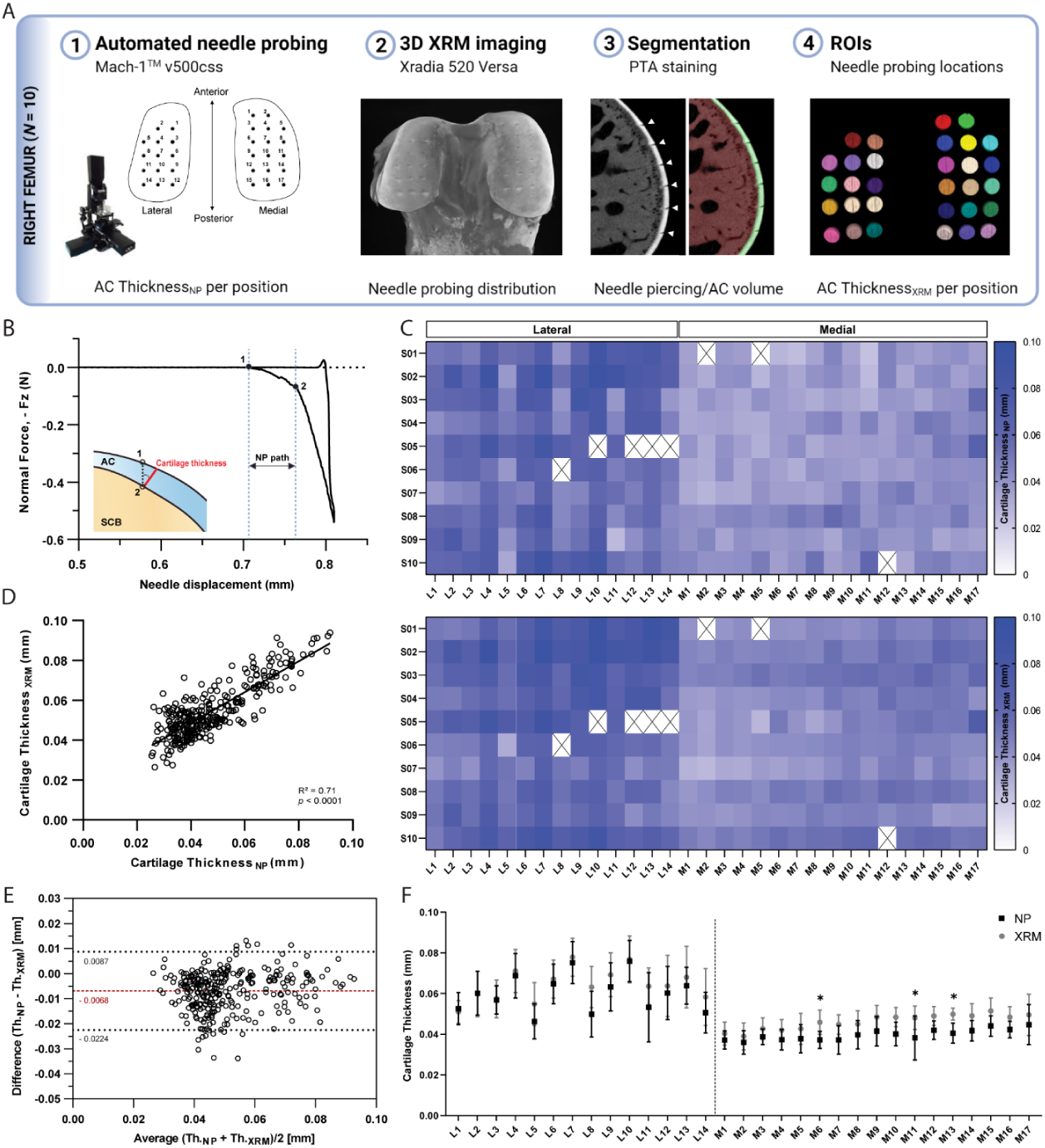
(A) Assessment of agreement between needle probing (NP) and XRM imaging cartilage thickness per position of measurement within right femoral condyles (*n =* 10) (B) Representative normal force-displacement curve obtained during NP test depicting articular cartilage, AC, surface (1) and subchondral bone, SCB, interface (2) positions and cartilage thickness calculated normal to the surface (*red)* using the surface angle orientation (C) Mapping distributions of cartilage thickness values per position as measured by NP and XRM (D) Correlation graph of cartilage thickness measured by NP vs. XRM, R = 0.842, n = 302, *p <* 0.0001 and corresponding (E) Bland-Altman plot showing overall agreement between methods, with average difference of 6.8 µm in thickness. Dotted black lines show upper and lower 95% limit of agreement F) Pairwise assessment of mean cartilage thickness NP vs. XRM per position for the lateral and medial condyles (* *p <* 0.05, ** *p <* 0.01, *two-way* ANOVA). Symbols represent the means and error bars the standard deviation.

Unlike peak force distributions, spatial heterogeneities in cartilage thickness were apparent among individual testing sites only on the lateral condyle (*p <* 0.0001), with averaged values ranging from 46 to 76 µm; whereas thickness distributions within the medial condyle was more uniform (*p =* 0.06), ranging from 36 to 45 µm. Nevertheless, no significant differences were observed within condyles when comparing their anteroposterior sub-regions; whereas the medial condyle was significantly thinner than its lateral counterpart, both in its anterior and posterior sub-regions (Tables 2 and S2). Together, cartilage thickness appears as a contributing factor but not the sole explanation to mechanical variations, which is also affected by differences in composition and morphology.

**Table 2.** Mean and standard deviation (SD) values for cartilage thickness as determined by needle probing for distal femur samples of murine articular cartilage (*n* = 10). Mean values compared between condyles (Lateral/Medial; unpaired, Student’s *t* test) and within sub-regions of condyles (Lat/Ant, Lat/Post, Med/Ant, Med/Post; *one-way* ANOVA, α = 0.05).

| Cartilage Thickness [$\mu\text{m}$] | | | | |
| --- | --- | --- | --- | --- |
| Condyle | Mean (SD) | Condyle | Mean (SD) | Lat - Med |
| Lateral | 60.3 (6.3) | Medial | 39.8 (2.9) | $p < 0.0001$ |
| Lat/Ant | 57.0 (5.9) | Med/Ant | 37.5 (3.8) | $p < 0.0001$ |
| Lat/Post | 62.3 (7.8) | Med/Post | 41.9 (3.0) | $p < 0.0001$ |
| Ant - Post | *ns* | Ant - Post | *ns* | - |

*ns*, no significant difference; -, comparison not applicable

As expected, pairwise comparisons for individual testing sites demonstrated a significant negative correlation between peak force and thickness measurements within lateral (R = −0.554, *n =* 135, *p* < 0.001, Figure 3A) and medial (R = −0.463, *n =* 167, *p* < 0.001, Figure 3B) condyles. Knowledge of site-specific thickness variations allowed the compressive stiffness to be determined for the same mechanical strain at each testing site. Instantaneous modulus was calculated using Hayes *et al*. elastic model ^31^ at 20% strain, wherein linear elastic behavior can be assumed and instantaneous response is considered as flow-independent (*Poisson’s* ratio, υ = 0.5 assumed) ^35^. Regardless of femoral condyle, instantaneous stiffness demonstrated no significant correlation to thickness variations (Figures 3C-D). Notably, compressive stiffness differed significantly between condyles (Lat vs. Med, *p <* 0.001), but no longer within a condyle (Lat/Ant vs Lat/Post: *p >* 0.99; Med/Ant vs Med/Post: *p* = 0.546), like seen for peak reaction force on the medial side. Next, we assessed the potential of this indentation testing in identifying microscale biomechanical differences between healthy and degraded cartilage.

**Figure 3.**
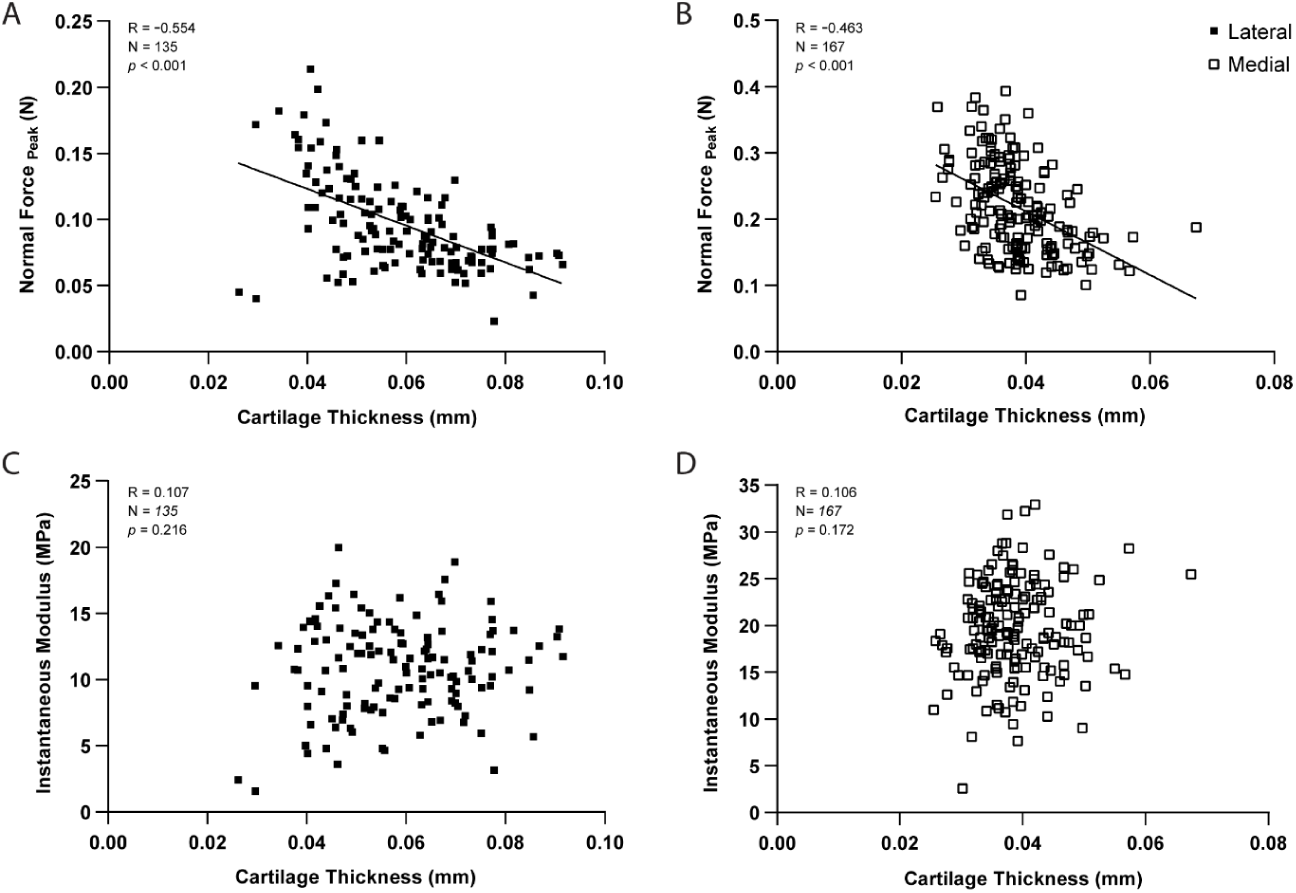
Correlation graphs per testing site for the lateral (*n* = 135 positions) and medial (*n =* 167) condyles, showing cartilage thickness (needle probing) is significantly correlated to peak indentation force at 20 um (A-B), but not to instantaneous modulus values as determined by Hayes *et al*. (1972) elastic model at 20% strain. *p-*values reported.

### Altered biomechanical properties in degenerated murine articular cartilage

To assess the changes in mechanical response within the context of cartilage degeneration, we employed the same testing protocol on age-matched Proteoglycan 4 (PRG4) knockout mice (*n* = 6) and compared the outcomes to the C57Bl/6 controls. PRG4 is a mucin-like glycoprotein highly conserved across species ^36,37^ and functionally relevant in joint homeostasis and lubrication ^33,38,39^. PRG4 loss of function, as seen in knockout mice (*Prg4*^*-/-*^), leads to degenerative joint changes recapitulating the phenotype of human camptodactyly-arthropathy-coxavara-pericarditis (CACP) syndrome ^33,40^. Histological alterations of articular cartilage have been comprehensively described, and include surface roughness, tissue thickening, and loss of collagen parallel orientation at the superficial layer, progressing to irreversible tissue damage with age ^33,41^. Yet, the microscale assessment of site-specific mechanical variations, aside from friction, has not been described so far for *Prg4*^*-/-*^ knee cartilage surfaces. Mapping of biomechanical parameters allowed site-specific differences in *Prg4*^-/-^ cartilage to be visualized, and outcomes were largely reproducible across femoral specimens (Figure 4A-B). These findings were supported by the combined quantitative assessment between genotypes for the different condyle regions and subregions (Figure 4C-F). *Prg4*^*-/-*^ mean cartilage thickness on both lateral (67.9 ± 3.5 µm, *p* = 0.032) and medial (56.3 ± 3.7 µm, *p <* 0.001) sides of the knee was higher compared to controls (Figure 4C). Differences in cartilage thickness between genotypes on anteroposterior sub-regions, however, were only detected on the medial condyle (Figure 4E). The site-specific thickness measurements enabled us to determine the instantaneous modulus at each testing site for the same mechanical strain. The utilized Hayes *et al*. elastic model could fit the response of mouse cartilage with great accuracy (RMSE equal to 0.017 ± 0.007 MPa for controls and 0.010 ± 0.003 MPa for *Prg4*^*-/-*^). Cartilage compressive stiffness was significantly lower on the medial condyle of *Prg4*^-/-^ mice compared to controls (8.67 ± 1.79 MPa vs. 19.52 ± 2.44 MPa, respectively, *p* < 0.001). Similar patterns were seen when considering the anteroposterior subregions of the medial condyle (Figures 4F). No significant differences in lateral condyle cartilage stiffness were observed (10.85 ± 1.73 MPa - control vs. 8.73 ± 2.39 MPa - *Prg4*^*-/-*^, *p=* 0.187).

**Figure 4.**
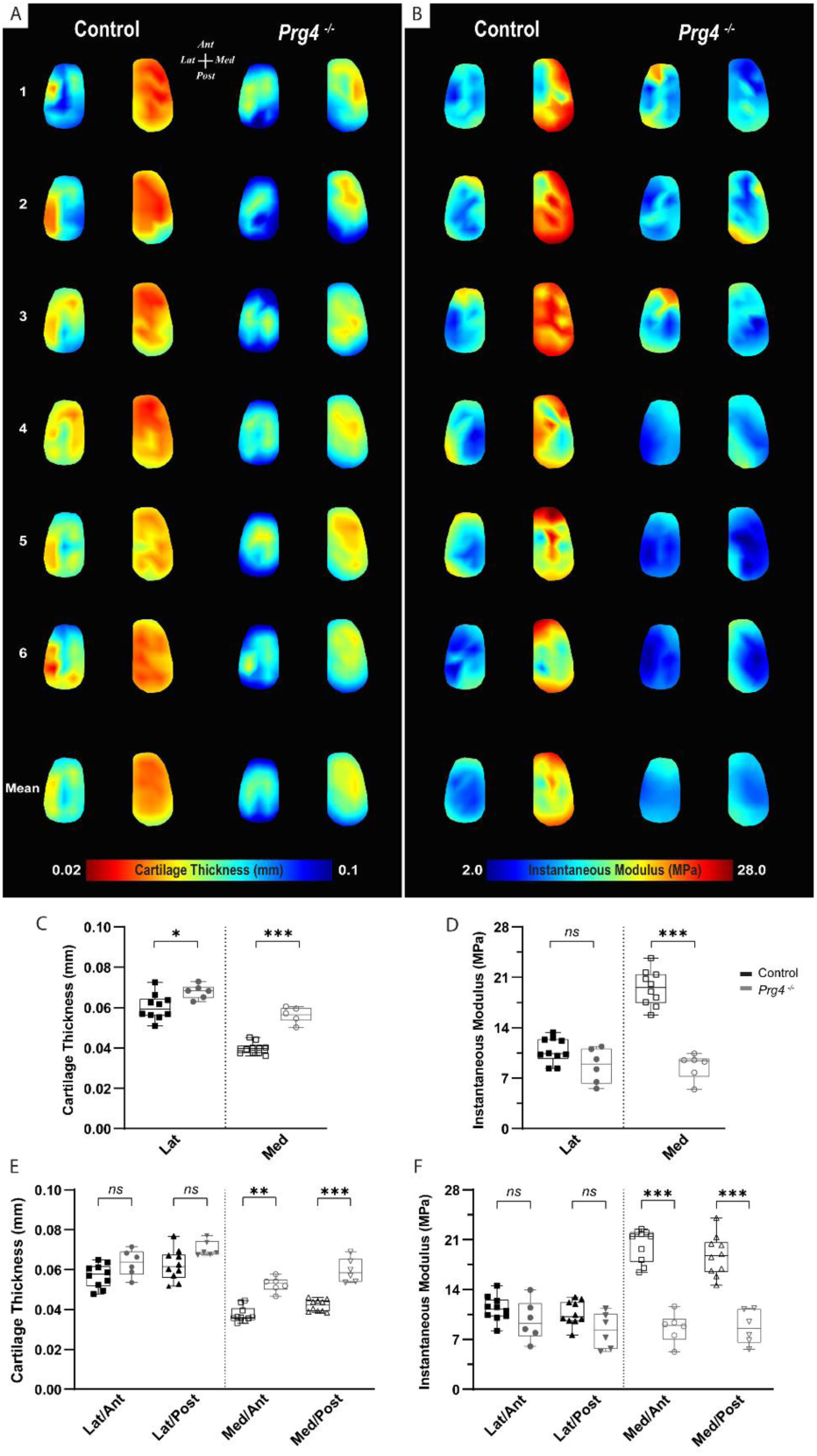
(A-B) Heat maps and corresponding boxplots comparing (C-D) regions - lateral and medial - and their (E-F) sub-regions - Lat/Ant, Lat/Post, Med/Ant, Med/Post - highlight spatial differences in biomechanical parameters on femoral cartilage surface between controls (C57Bl/6, *n =* 10) and PRG4 knockout mice (*Prg4*^*-/-*,^ *n =* 6). Maps shown for six representative samples per genotype, as well as the corresponding averaged map of all samples. Parameters illustrated are thickness measured by NP (A, C, E) and instantaneous modulus as determined by Hayes *et al*. (1972) elastic model (B, D, F). Pairwise comparison between genotypes for mean values on the lateral and medial condyles and anteroposterior sub-regions (*Mann-Whitney U* with *Bonferroni-Dunn* correction, ^*^*p* < 0.05, ^**^*p* < 0.01, ^***^*p* < 0.001).

## Discussion

Using a novel microscale instrumented apparatus, we were able to reliably detect and quantify spatial variations in biomechanical parameters across murine cartilage surfaces, both within healthy and degenerated femoral condyles. Compared to recent mouse studies using this commercially available apparatus ^26,42^, the optimization of sample mounting and needle probing protocols allowed for unprecedented quantitative mapping of the mechanical behavior and associated cartilage thickness on load-bearing regions of distal femurs with high spatial density.

For healthy, 4-month old C57Bl6 mice, micro-indentation measurements were reproducible at any given testing site, also indicating that the instantaneous deformation employed and subsequently sustained compression protocol did not compromise the mechanical behavior of the freshly harvested cartilage tissue. In addition, heterogeneity in peak forces were identified across anatomical locations. Notably, the medial condyle surface yielded approximately two times higher peak forces on average than its lateral counterpart. Moreover, both condyles displayed a shift towards lower values at posterior regions. Previous studies have shown significant variation in mechanical properties within different cartilage surfaces in healthy joints ^43,44^, or even over a single surface ^18,45^. However, the effect of site-specific cartilage geometry, such as thickness, when characterizing the mechanical properties of cartilage in indentation is often overlooked ^23,46^.

Studies in mice commonly rely on mean thickness values from histology, requiring longer turnaround and lacking spatial specificity. The optimized needle probing protocol used here, was able to resolve spatial thickness variations across condyle surfaces, and for the first-time closely localize cartilage thickness measures to the footprint of indentation on mouse distal femurs. Moreover, three-dimensional visualization of the femoral cartilage surfaces after XRM imaging validated the positioning and distribution of the testing grid used, allowed us to pinpoint individual testing sites and compute their thickness. A highly significant correlation between pairwise cartilage thickness values from NP and XRM, per measurement site, was observed. Small variations seen between averaged thickness values could be explained by differences in instrument resolution capabilities (Mach-1 z-axis: 0.5 µm vs XRM: 4.39 µm voxel size), partial volume effects in XRM segmentation, and/or compromise of cartilage integrity by needle probing, which might have led to cartilage swelling and overestimation on subsequent XRM thickness measures. Nevertheless, mean thickness values were in line with earlier reports for mouse cartilage ^29,47,48^, and marked greater cartilage thicknesses were measured on the lateral condyle compared to the medial counterpart ^12^.

In general, higher peak reaction forces in intact cartilage were associated with lower thickness values at the same indentation depth, and likely amplified by contribution of the underlying subchondral bone on thinner areas ^49^. Localized cartilage thickness measurements enabled us to account for the effect of cartilage geometry when calculating compressive stiffness distributions. Quantitative analysis was adjusted to 20% strain (considered as physiological loading) ^50^, and corresponding instantaneous modulus could be derived from Hayes’s analytical formulation. According to the present results, there was no significant variation in mean thickness of healthy cartilage over a condyle surface, therefore biomechanical parameters were compared between matched regions and sub-regions when assessing degeneration.

Compared to C57Bl6 control, the loss of *Prg4* function resulted in an increase in thickness and decrease in stiffness of the knee cartilage in 16-weeks-old mice. This is consistent with previous observations on hip articular cartilage of *Prg4*^*-/-*^ mice (same age), using nanoscale atomic force microscopy (AFM) indentation ^39^. The cartilage thickness values determined by needle probing in our study are in a similar range to those reported by Karamchedu *et al*. ^41^. In their microCT imaging study, *Prg4*^*-/-*^ mice displayed cartilage thickness averaging 61.0 ± 4.3 µm and 42.4 ± 2.6 µm for the load-bearing regions of the lateral and medial condyle, respectively. Furthermore, *Prg4*^*-/-*^ cartilage was significantly thicker than in control littermates, even though mice were younger (10-weeks-old) than in our study.

Reduction in the instantaneous compressive modulus in our study was confined to the medial compartment. We attribute that to differences in thickening in *Prg4*^*-/-*^ femoral cartilage compared to controls, 12.6% versus 41% on lateral and medial condyles respectively, as well as *Prg4*^*-/-*^ compromised structural integrity ^33,38,51^ having a bigger functional impact on the medial compartment, recognized as the main load bearing region in most mammalian joints. Microscale indentation characterizes the overall tissue resistance to deformation ^50^. Due to the rapid compression rate employed in the present protocol, normal cartilage tissue is expected to deform with minimal change in volume ^35^, as its low permeability restrains fluid flow; thereby localizing the strain near the surface as fluid pressure pushes against the collagen meshwork. Thus, under instantaneous (impact) loading the ability of cartilage to resist compression is known to be affected by the collagen fibril meshwork ^49,52^, particularly tangentially oriented collagen fibrils on the superficial tissue layer ^43^. In *Prg4*^-/-^ the normal parallel organization of collagen fibrils adjacent to the surface is known to be disrupted ^38^, likely affecting tissue permeability as well, helping explain the lower instantaneous modulus and slightly faster dissipation of stress.

Limitations of this study include evaluation of the anatomy of the distal murine femoral condyles and not the opposing articulation (i.e., proximal tibia). Also, we focused on a single timepoint analysis, however, investigations related to aging and progressive degeneration are of interest, as they are known to affect the biochemical composition and structural integrity of cartilage ^33,53,54^, thereby influencing its mechanical properties. Finally, due to the biphasic/poroviscoelastic nature of cartilage, future studies could consider the use of more complex analytical models ^55,56^ able to capture the viscoelastic behavior of cartilage in mice.

In this study, we have gained insights into the patterns of varying surface geometry, and mechanics present within murine articular cartilage at the microscale. Three-dimensional indentation mapping was able to resolve site-specific differences in thickness and mechanical properties across knee cartilage surfaces in healthy mice. Moreover, it identified functional changes on the *Prg4*^*-/-*^ mouse model. This technique could also prove helpful for the study of other mouse models mimicking different mechanisms of, or therapies focused on, repair and degeneration of articular cartilage, as microscale indentation with high spatial density can provide a more comprehensive characterization of cartilage’s mechanical properties.

## Materials and methods

### Ethics statement

All mouse experiments were carried out following the Canadian Council on Animal Care Guidelines recommendations and approved by the University of Calgary Animal Care Committee (protocols AC16-0043 and AC20-0042).

### Animals

Male C57Bl6 mice were purchased from *Jackson Laboratories* (Bar Harbor, ME). We based on sample on a recent paper that used indentation (but not needle probing) in mice ^26^, however, because of the addition of needle probing we decided to increase the sample size. Animals were housed under a standard light cycle and had free access to feed (standard diet) and water. Ten mice were euthanized at 16-weeks of age, and hind limbs (*n* = 10 right) were harvested for biomechanical testing and 3D X-ray microscopy (XRM) imaging. Age-matched PRG4 knockout mice (*Prg4*^*-/-*^, *n* = 6) were generated and maintained on a C57BL/6 genetic background, as previously described ^57^. Limbs were disarticulated at the hip, followed by transection of the ligaments and careful isolation of distal femurs from tibiae and menisci with the help of a dissection microscope (*Leica*). Femurs were preserved gently wrapped in Kimwipe soaked in phosphate-buffered saline (PBS, pH 7.4) until the time of assessment. All samples were mechanically tested no longer than 3h after dissection to prevent tissue degradation.

### Automated Indentation Mapping

The shafts of isolated femurs were glued into a 0.1-10 µL pipette tip (*VWR*) using cyanoacrylate adhesive, fixed into a stainless-steel hex nut (*Paulin*, Model 848-216) and secured to the sample holder (Figure S1). This customized setup allowed for simple and proper positioning of the sample exposing the load-bearing region of the condyles ^27^ for data acquisition, as well as non-destructive retrieval of samples after testing, such that subsequent XRM imaging could be carried out. A standardized mapping grid (*n* = 31 positions) was superimposed on an image of the cartilage surface, consisting of 14 and 17 measurement sites at the lateral and medial condyles, respectively (Figure 1A). The testing chamber was filled with PBS solution at room temperature, and the tissue was allowed to equilibrate before testing. Automated indentation mapping under stress-relaxation was then performed using the Mach-1TM v500css (Biomomentum Inc., Laval, QC) device, equipped with a calibrated multiple-axis load cell (±17 N, 3.5 mN force resolution) and associated software. With each site, the contact coordinates (and contact coordinates at four adjacent places in a 0.075 mm scanning grid) were measured using a contact criteria of 0.035 N. Surface orientation was then calculated using the surrounding contact coordinates. Indentation was achieved by moving the indenter perpendicular to the surface while moving the Mach-1 stages concurrently in all 3-axis. The spherical indenter (0.3 mm in diameter) was driven into the cartilage to a depth of 20 μm over one second and held constant for 90 seconds. For C57Bl6, a total of 3 indentation tests were performed per sample, approximately 45 minutes apart. Data reported consist of peak force and instantaneous modulus, as determined by fitting the Hayes et al. (1972) ^31^ elastic model to the load-displacement curves at 20% strain. Assessment of how well the model fit the resulting curve per test site was done using root mean square error (RSME). Since the analysis per position across specimens and genotypes reflect deviations due to anatomical positioning, calculated parameters were compared between lateral (Lat) and medial (Med) femoral condyles, as well as on four condylar sub-regions (Lateral/Anterior - Lat/Ant, Lateral/Posterior – Lat/Post, Medial/Anterior - Med/Ant, Medial/Posterior - Med/Post), each containing at least 5 positions of measurement (Figure 1A).

### Needle probing – thickness measurement

After indentation mapping, the spherical indenter was replaced by a 30G x 1.4” hypodermic needle (TSK Laboratory, Japan) adapted to the 1 mm spherical indenter (*Biomomentum Inc*., Laval, Canada) (Figure S2). Automated thickness mapping (Figure 2A) was done on the knee cartilage surface using the needle probing technique ^32^, and previous mapping grid distribution slightly shifted as appropriate to account for the off-axis position of the needle’s bevel tip. The needle was driven vertically into the cartilage surface at a constant speed until a 0.5 N stop criteria was reached in the subchondral bone. The cartilage surface and cartilage/subchondral bone interface positions were identified in the load-displacement curves (Figure 2B) generated at each measurement site using the automatic mode of analysis ^24^. A 0.25 N/s loading limit was defined to identify the interface position. Manual correction was employed when the algorithm failed to identify the inflection point ^24^. The cartilage thickness reported corresponds to the vertical needle displacement from cartilage surface to subchondral bone multiplied by the cosine of the surface orientation angle (Figure 2B) as determined during automated indentation for each position. After testing, samples were preserved in 10% NBF solution for 24h and stored in 70% ethanol for 24h to 48h before imaging.

### 3D X-ray Microscopy (XRM) imaging

3D XRM imaging was used for non-destructive assessment of cartilage morphology after biomechanical testing. Fixed femurs were incubated for 16-18h in 1% phosphotungstic acid (PTA) solution at room temperature for cartilage contrast enhancement before imaging ^15^. Samples were enclosed onto a custom specimen chamber, with 1% PTA in 70% ethanol added to the chamber’s bottom to minimize tissue dehydration. Zeiss Xradia 520 versa (*Carl Zeiss* X-Ray Microscopy, Pleasanton, CA) scans of each distal femur were obtained following previously described protocol ^58,59^. In brief, high-resolution scans of 2001 axial slices were acquired at a 4.39 µm voxel size, with low-energy (40 kVp voltage, 3 W power) x-rays.

### Imaging processing

The contrast-enhanced cartilage surface was segmented by determining a threshold intensity, thereby delineating the femur scan into cartilage and subchondral bone voxels. Needle probing left a physical deformity in the articular cartilage of the right femurs visible on XRM imaging (Figure 2A), allowing for all 31 regions of interest (ROIs) corresponding to needle probing to be manually landmarked. Landmarks were placed manually using the two-dimensional axial, sagittal, and coronal planes centered along the cartilage thickness and within each needle probing site. Cartilage segmentations were corrected manually to ensure the cartilage mask encompassed resulting volume gaps at needle probing positions. Then, the segmented cartilage was masked by a sphere of radius 75 µm placed on each landmark, leaving a thin disk of cartilage. The thickness transform was computed for each disk ^60^ and the mean thickness values, taken as a statistic of the thickness distribution, were used to minimize variability due to morphological changes in the cartilage caused by the mechanical testing. Image processing was performed in SimpleITK ^61^ (Insight Software Consortium, v1.2.4), and morphometry was performed in Image Processing Language (IPL v5.42, SCANCO Medical AG, Brüttisellen, Switzerland).

### Statistical analysis

Analyses were performed in GraphPad Prism software (version 9), *α* = 0.05 was considered statically significant. Continuous parameters are reported as mean values and corresponding standard deviations (SDs). Normality was assessed by Shapiro-Wilk normality test. C57Bl6 peak load and thickness data was analysed by Student *t* test or ANOVA, with Bonferroni *post-hoc* comparisons. To assess differences in biomechanical properties between genotypes non-parametric Mann-Whitney U test with *Bonferroni-Dunn’s* correction was used. To test reliability and absolute agreement between repeated measurements, SPSS 27 (*IBM*, Chicago, IL) was used to obtain single-measurement, two-way mixed effect intraclass correlation coefficient estimates and respective 95% limits of agreement ^62,63^. When assessing cartilage thickness measurements between needle probing and XRM imaging techniques, *Pearson* correlation coefficient (R) was used for method-comparison and Bland-Altman analysis to assess bias between methods.

## Acknowledgments

Graphical representation of experimental designs was created using BioRender.com. The authors thank *Biomomentum inc*. for their assistance during optimization of indentation testing protocol. AOM would like to thank the University of Calgary and Alberta Innovates for support during the period this study was conducted.

## Author’s contribution

A.O.M. was responsible for the study design, data collection, data analysis, data interpretation and draft of the manuscript. B.B. carried out imaging processing and data analysis. W.B.E. contributed to the study design. R.K. contributed to the study design and data interpretation. B.B., W.B.E. and R.K. provided critical review, commentary and revisions to the manuscript.

## Supplementary Material

**Figure S1.**
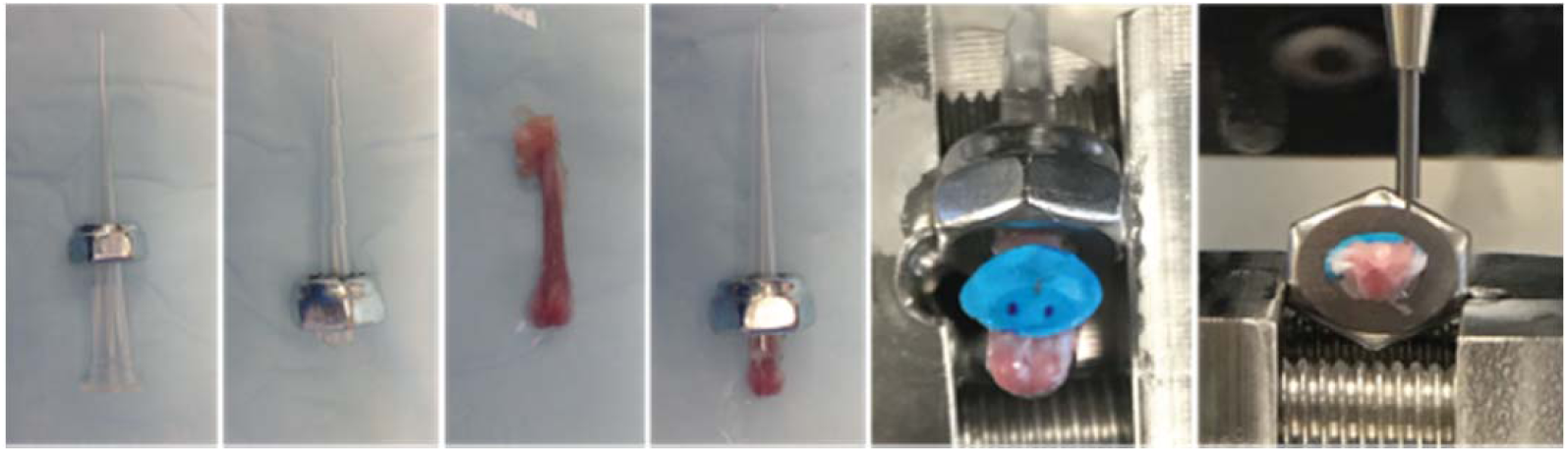
Specimen preparation and assemble to sample holder for automated indentation mapping using custom setup, allowing for repositioning of the sample and non-destructive retrieval for 3D XRM imaging.

**Table S1.** Mean peak force 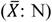 and coefficient of variation (CV: %) from triplicate measurements for each of the 31 positions (L1-M17) of assessment over mice femoral condyles.

|  | S01 |  | S02 |  | S03 |  | S04 |  | S05 |  | S06 |  | S07 |  | S08 |  | S09 |  | S10 |  |
| --- | --- | --- | --- | --- | --- | --- | --- | --- | --- | --- | --- | --- | --- | --- | --- | --- | --- | --- | --- | --- |
| | $\bar{X}$ | CV | $\bar{X}$ | CV | $\bar{X}$ | CV | $\bar{X}$ | CV | $\bar{X}$ | CV | $\bar{X}$ | CV | $\bar{X}$ | CV | $\bar{X}$ | CV | $\bar{X}$ | CV | $\bar{X}$ | CV |
| L1 | 0.08 | 2.9 | 0.08 | 3.0 | 0.10 | 4.1 | 0.13 | 1.8 | 0.09 | 12.1 | 0.10 | 2.7 | 0.10 | 4.9 | 0.09 | 3.3 | 0.07 | 1.5 | 0.11 | 0.2 |
| L2 | 0.11 | 1.3 | 0.09 | 2.5 | 0.10 | 2.3 | 0.14 | 6.1 | 0.12 | 8.2 | 0.11 | 4.5 | 0.13 | 4.9 | 0.11 | 6.1 | 0.07 | 13.7 | 0.11 | 5.7 |
| L3 | 0.07 | 4.9 | 0.08 | 3.6 | 0.08 | 5.0 | 0.12 | 2.5 | 0.11 | 4.8 | 0.12 | 1.5 | 0.09 | 3.8 | 0.08 | 4.9 | 0.07 | 8.5 | 0.08 | 8.0 |
| L4 | 0.10 | 7.2 | 0.07 | 2.8 | 0.10 | 1.3 | 0.11 | 1.6 | 0.08 | 3.6 | 0.11 | 4.7 | 0.13 | 2.2 | 0.09 | 3.0 | 0.06 | 4.8 | 0.09 | 5.5 |
| L5 | 0.16 | 0.7 | 0.15 | 5.0 | 0.21 | 9.0 | 0.14 | 7.7 | 0.12 | 7.8 | 0.04 | 10.0 | 0.17 | 2.4 | 0.17 | 4.2 | 0.11 | 6.0 | 0.19 | 6.3 |
| L6 | 0.06 | 9.3 | 0.07 | 7.4 | 0.08 | 13.4 | 0.12 | 4.1 | 0.09 | 5.7 | 0.08 | 2.9 | 0.06 | 9.0 | 0.06 | 6.9 | 0.08 | 6.0 | 0.07 | 5.8 |
| L7 | 0.06 | 4.7 | 0.07 | 6.2 | 0.08 | 6.9 | 0.10 | 3.1 | 0.07 | 12.3 | 0.07 | 3.8 | 0.09 | 8.0 | 0.08 | 4.9 | 0.07 | 3.8 | 0.07 | 7.5 |
| L8 | 0.16 | 2.2 | 0.09 | 15.9 | 0.19 | 3.2 | 0.14 | 4.9 | 0.11 | 7.9 | - | - | 0.20 | 5.8 | 0.17 | 3.1 | 0.12 | 3.5 | 0.12 | 6.5 |
| L9 | 0.05 | 10.3 | 0.07 | 5.4 | 0.08 | 2.4 | 0.09 | 1.5 | 0.03 | 20.2 | 0.07 | 0.3 | 0.05 | 6.4 | 0.07 | 2.9 | 0.09 | 0.9 | 0.08 | 2.5 |
| L10 | 0.04 | 4.2 | 0.07 | 6.8 | 0.07 | 3.1 | 0.08 | 4.5 | - | - | 0.08 | 3.3 | 0.07 | 3.9 | 0.07 | 5.5 | 0.07 | 1.1 | 0.08 | 3.2 |
| L11 | 0.10 | 5.7 | 0.08 | 6.1 | 0.15 | 2.4 | 0.11 | 9.1 | 0.13 | 5.1 | 0.14 | 6.8 | 0.16 | 5.9 | 0.14 | 3.7 | 0.04 | 9.0 | 0.11 | 1.3 |
| L12 | 0.05 | 0.9 | 0.08 | 4.6 | 0.07 | 9.4 | 0.07 | 3.6 | - | - | 0.09 | 3.1 | 0.06 | 6.2 | 0.06 | 8.8 | 0.11 | 3.6 | 0.10 | 3.3 |
| L13 | 0.05 | 3.0 | 0.07 | 3.6 | 0.07 | 6.4 | 0.07 | 5.8 | - | - | 0.10 | 2.3 | 0.07 | 2.9 | 0.08 | 4.9 | 0.09 | 0.9 | 0.09 | 6.0 |
| L14 | 0.08 | 5.0 | 0.08 | 5.7 | 0.13 | 4.9 | 0.10 | 4.9 | 0.04 | 19.0 | 0.14 | 1.1 | 0.14 | 3.7 | 0.16 | 1.7 | 0.09 | 12.0 | 0.12 | 1.4 |
| M1 | 0.29 | 3.7 | 0.18 | 6.9 | 0.23 | 4.1 | 0.23 | 1.9 | 0.21 | 5.5 | 0.26 | 1.4 | 0.22 | 6.7 | 0.20 | 2.2 | 0.20 | 1.7 | 0.22 | 1.9 |
| M2 | - | - | 0.25 | 4.6 | 0.25 | 3.2 | 0.26 | 1.0 | 0.28 | 2.3 | 0.29 | 2.1 | 0.27 | 1.5 | 0.24 | 6.3 | 0.20 | 5.4 | 0.29 | 0.9 |
| M3 | 0.23 | 3.1 | 0.17 | 3.8 | 0.24 | 0.7 | 0.24 | 1.4 | 0.24 | 2.8 | 0.24 | 1.8 | 0.23 | 3.2 | 0.19 | 2.7 | 0.22 | 6.7 | 0.25 | 3.4 |
| M4 | 0.36 | 5.0 | 0.30 | 6.5 | 0.28 | 0.6 | 0.30 | 2.6 | 0.29 | 3.1 | 0.31 | 2.7 | 0.25 | 5.8 | 0.26 | 3.1 | 0.19 | 4.5 | 0.30 | 2.7 |
| M5 | - | - | 0.38 | 10.0 | 0.30 | 3.9 | 0.30 | 1.9 | 0.36 | 1.1 | 0.26 | 0.7 | 0.30 | 2.4 | 0.25 | 5.5 | 0.18 | 4.3 | 0.15 | 5.1 |
| M6 | 0.19 | 1.7 | 0.17 | 6.3 | 0.25 | 3.4 | 0.23 | 3.8 | 0.19 | 5.6 | 0.26 | 2.3 | 0.21 | 0.7 | 0.11 | 10.9 | 0.16 | 3.9 | 0.20 | 15.1 |
| M7 | 0.30 | 3.3 | 0.30 | 4.2 | 0.34 | 5.9 | 0.30 | 1.9 | 0.37 | 7.0 | 0.31 | 1.7 | 0.26 | 2.2 | 0.28 | 1.1 | 0.16 | 1.4 | 0.30 | 6.9 |
| M8 | 0.37 | 3.3 | 0.36 | 6.2 | 0.33 | 2.0 | 0.28 | 5.1 | 0.35 | 5.6 | 0.23 | 2.0 | 0.23 | 6.0 | 0.17 | 3.6 | 0.13 | 1.7 | 0.14 | 4.2 |
| M9 | 0.20 | 3.8 | 0.16 | 2.5 | 0.27 | 5.6 | 0.21 | 2.8 | 0.18 | 19.6 | 0.25 | 1.2 | 0.20 | 4.2 | 0.16 | 0.9 | 0.15 | 3.4 | 0.24 | 18.6 |
| M10 | 0.26 | 2.8 | 0.30 | 2.0 | 0.33 | 7.1 | 0.28 | 1.2 | 0.39 | 2.2 | 0.25 | 2.4 | 0.19 | 2.5 | 0.25 | 2.0 | 0.16 | 2.4 | 0.24 | 2.7 |
| M11 | 0.28 | 2.0 | 0.30 | 2.8 | 0.30 | 6.0 | 0.21 | 1.6 | 0.31 | 3.8 | 0.22 | 2.0 | 0.16 | 1.1 | 0.14 | 0.7 | 0.13 | 7.8 | 0.11 | 7.1 |
| M12 | 0.16 | 1.6 | 0.19 | 5.9 | 0.25 | 1.6 | 0.18 | 0.6 | 0.17 | 2.3 | 0.18 | 6.4 | 0.15 | 6.9 | 0.18 | 5.7 | 0.09 | 7.7 | - | - |
| M13 | 0.22 | 4.3 | 0.26 | 4.6 | 0.29 | 3.0 | 0.20 | 2.3 | 0.32 | 4.0 | 0.20 | 2.6 | 0.15 | 1.2 | 0.21 | 3.6 | 0.14 | 0.1 | 0.20 | 2.9 |
| M14 | 0.19 | 1.6 | 0.27 | 1.2 | 0.25 | 4.2 | 0.17 | 2.6 | 0.27 | 1.2 | 0.20 | 0.8 | 0.14 | 2.9 | 0.13 | 4.1 | 0.12 | 3.2 | 0.12 | 1.8 |
| M15 | 0.16 | 3.7 | 0.16 | 0.5 | 0.18 | 1.6 | 0.14 | 1.6 | 0.18 | 0.7 | 0.13 | 1.9 | 0.13 | 1.7 | 0.14 | 1.1 | 0.14 | 1.6 | 0.21 | 9.2 |
| M16 | 0.14 | 1.3 | 0.19 | 1.3 | 0.20 | 0.9 | 0.14 | 2.1 | 0.22 | 1.4 | 0.16 | 2.7 | 0.12 | 0.8 | 0.15 | 1.2 | 0.13 | 1.4 | 0.16 | 1.7 |
| M17 | 0.14 | 2.2 | 0.21 | 1.1 | 0.17 | 3.4 | 0.15 | 2.6 | 0.19 | 3.1 | 0.15 | 5.5 | 0.13 | 2.1 | 0.12 | 3.0 | 0.13 | 1.4 | 0.13 | 2.2 |

**Figure S2.**
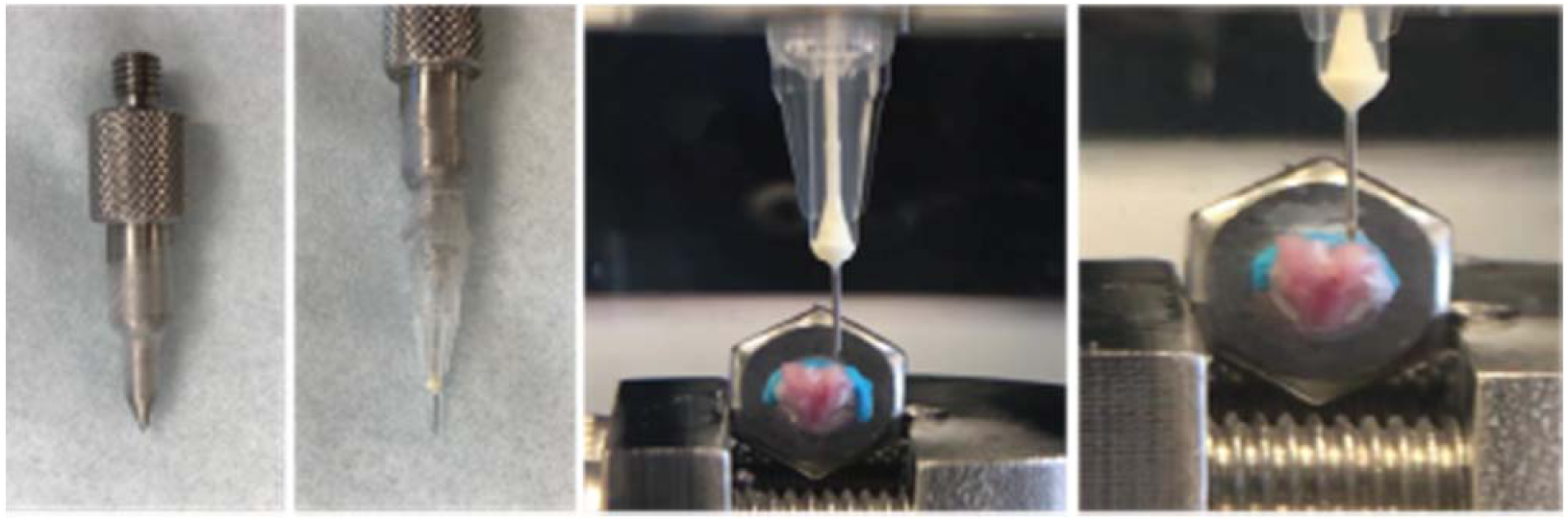
Setup for needle probing thickness measurement, using a 30G x 1.4” hypodermic needle (TSK Laboratory, Japan) adapted to the 1 mm spherical indenter (*Biomomentum Inc*., Laval, QC).

**Table S2.** Mean and standard deviation (SD) values for cartilage thickness as determined by XRM imaging for distal femur samples of murine articular cartilage (*n* = 10). Mean values compared between condyles (Lateral/Medial; unpaired, Student’s *t* test) and within sub-regions of condyles (Lat/Ant, Lat/Post, Med/Ant, Med/Post; *one-way* ANOVA). *p*-value reported.

| Cartilage Thickness [N] |  |  |  |  |
| --- | --- | --- | --- | --- |
| Condyle | Mean (SD) | Condyle | Mean (SD) | Lat - Med |
| Lateral | 64.4 (8.4) | Medial | 46.5 (4.4) | $p < 0.0001$ |
| *Lat/Ant* | 59.0 (8.2) | *Med/Ant* | 42.9 (5.1) | $p < 0.0001$ |
| *Lat/Post* | 67.4 (9.7) | *Med/Post* | 49.5 (4.3) | $p < 0.0001$ |
| Ant - Post | *ns* | Ant - Post | *ns* | - |

*ns*, no significant difference; -, comparison not applicable

